# Intrinsic and Extrinsic Modulation of C. elegans Locomotion

**DOI:** 10.1101/312132

**Authors:** Jack E. Denham, Thomas Ranner, Netta Cohen

**Affiliations:** School of Computing, University of Leeds, UK

**Keywords:** proprioception, neural control, biomechanics, undulatory locomotion, nematodes

## Abstract

Animal locomotion describes the coordinated self-propelled movement of a body, subject to the combined effects of internal muscle forcing and external forces. Here we use an integrated neuromechanical computational model to study the combined effects of neural modulation, mechanical modulation and modulation of the external environments on undulatory forward locomotion in the nematode *C. elegans*. In particular we use a proprioceptively driven neural control circuit to consider the effects of proprioception, body elasticity and environmental drag on the waveform, frequency and speed of undulations. We find qualitative differences in the frequency-wavelength relationship obtained under extrinsic modulation of the environmental fluid or body elasticity versus intrinsic modulation due to changes in the sensorimotor control. We consider possible targets of modulation by the worm and implications of our results for our understanding of the neural control of locomotion in this system.

## INTRODUCTION

Undulations, or movement via the propagation of mechanical waves along a body, is a remarkably old and successful strategy for locomotion, and one that is prevalent across all scales of life – from microorganisms to monster prehistoric snakes.^8^ While each life form is unique, the generation of whole body undulations resulting in directional movement necessarily emerges from the coupling among the components of the system, including the nervous system and muscles in animals, other body tissue, and the physical environment. Understanding the separate and combined roles that these components play can help elucidate the constraints imposed on the neuromechanical system and any targets for internal modulation. The small, compact anatomy and fully mapped nervous system of the nematode *Caenorhabditis elegans* along with its undulatory repertoire make it an ideal organism for linking neural control with behavior.^30,31^ Like other organisms,^15^ *C. elegans* exhibits gait modulation when swimming through media characterized by different viscosities or viscoelasticities. Higher resistance due to external drag forces results in slower undulations with shorter wavelength and lower wave amplitude.^1,10^ While internal neural control can also affect speed and waveform within homogeneous environments,^6,7,21,27^ the coupling between internal and external modulation of gait has received little attention. This question is particularly interesting in systems where proprioception – the sensing of the position or movement of different parts of the body – plays a crucial role in the generation or entrainment of the motor patterns, as is generally accepted in *C. elegans*.^4,5,10,12,13,16,22,29^

We use an integrated neuromechanical computational model to address this question. The model combines biomechanical realism, seamless integration of biomechanics with neuromuscular control, numerical stability and high computational efficiency, allowing us to study a range of hypotheses and perform systematic parameter sweeps (430 simulations in this paper) to gain understanding and mechanistic insight about both the neural control and neuromechanical coupling.

The neural circuit explored here lacks central pattern generation and instead relies on proprioceptive feedback to generate undulations. In what follows, we first show that this model is capable of replicating the swim-crawl transition, previously observed experimentally in different concentrations of media^1,12^ and theoretically captured by Boyle *et al*. in an articulated neuromechanical framework with similar proprioceptive control.^3^ We then use this framework to analyze the combined effects of the drag coefficients of the environment, the elasticity of the body and the effective proprioceptive control parameters in the motor circuit.

## MODEL OVERVIEW

Our model worm consists of a continuous incompressible viscoelastic shell, ^9^ with a formulation akin to viscoelastic filaments.^16^ At each point along the worm, we assume the width of the worm’s body is fixed in time, which allows us to collapse all internal (neuromuscular) and external (drag) forces onto the worm midline. We further impose a fixed length of 1 mm (along the midline). Four free parameters modulate the body properties and its interaction with the environment.

Environmental forces are modeled by resistive force theory and parametrized by two drag coefficients *K_ν_* and *K_τ_* that act to resist motion normal (*ν*) and tangential (*τ*) to the local body surface, respectively.^1,4,20^ For Newtonian fluids, including water (or buffer solution), we approximate the ratio of these drag coefficients at 1.5. Here we model both Newtonian and linear viscoelastic media, adopting existing estimates of drag coefficients for water^20^ 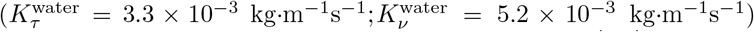 and for agar^1,4,22,28^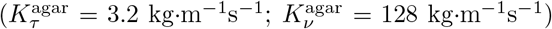, and interpolating coefficients for intermediate environments.

The passive body is parametrized by a Young’s modulus (the resistance to bending) and its internal viscosity (the body damping in response to bending). A nondimensionalization formulation shows that internal viscosity of the body may be neglected in sufficiently stiff external environments, such as agar. On the one hand, this implies that we restrict the validity of our simulations, but on the other, it offers us a better understanding of the limits of elastic models. Following previous analysis,^9,11^ we choose the default Young’s modulus to be 10^5^ Pa-high enough to allow the generation of thrust at reasonable speeds (on agar like environments). Note that the worm modulates its muscle stiffness (or elasticity) as a function of its activity.^24^ We assume that at any point in time, opposite muscles along the body contract and relax in antiphase, and further, for parsimony, that the sum of Young’s moduli on the two sides of the body is approximately constant. Thus, our choice Young’s modulus is an effective elasticity associated with the mean Young’s modulus of the body during undulatory behavior.

Muscles drive the above mechanical model with an active moment. They are modeled by a single leaky integration equation that converts a current input to a mechanical torque. The muscle forcing equation is associated with a single free parameter representing the muscle time constant, which we have fixed at 100 ms. As described in the accompanying paper,^11^ our neural control consists of a binary activation function acting continuously along the body, loosely representing ON/OFF neuromuscular activation by B-type excitatory motor neurons. At every point along the body, this switching is mediated by a threshold over the instantaneous proprioceptive input, integrated over its receptive range, posteriorly to the muscle coordinate. In the model, we use the simplest form of such proprioception to alternate ventral and dorsal activations, a single switch with only two free parameters: the proprioceptive range and the proprioceptive threshold.^11^ The latter is formulated as a threshold average curvature (in units of inverse length) over the proprioceptive range. Thus, in this model, proprioceptive neurons control the muscles, which generate an active moment driving a viscoelastic shell. The model equations balance internal and external forces and torques subject to mass conservation within the worm’s body. The mechanical model equations (except the neuromuscular generated active moment) and numerical (finite element) methods for their solution are given in full in Cohen and Ranner.^9^

## A CONTINUUM PROPRIOCEPTIVELY DRIVEN ELASTIC SHELL MODEL CAPTURES THE SWIM CRAWL TRANSITION

Before we use our model to investigate the effects of intrinsic modulations, we first asked whether this model successfully captures the spectrum of waveforms exhibited by the worm in different homogeneous media. Boyle *et al*.^3^ required some damping (or internal viscosity) to reproduce the observed gait modulation, particularly in environments close to water. In addition, that model also required an extended posteriorly facing proprioceptive range spanning half of the worm’s body length. We therefore initially picked the same proprioceptive range, and set the proprioceptive threshold to match the desired peak curvatures obtained on agar (10mm^−1^). To test whether, and under what constraints, this model captures gait modulation, we ran a large number of simulations, varying only the external drag coefficients, to mimic water-like, agar-like and intermediate environments.^1,4^ Some simulations maintained the ratio of drag coefficients fixed at 1.5, corresponding to Newtonian media, whereas others spanned a ratio of coefficients from 1.5 to 40.

Fig. 1 summarizes the gait modulation results obtained with our model. The qualitative trend confirms that with the chosen proprioceptive drive and model parameters, frequency and wavelength are tightly coupled. This choice of parameters also yields a quantitatively similar relationship between frequency and wavelength of undulations to those observed experimentally^1^. As we might expect, some frequencies are unrealistically high as we approach water so results in these environments should be interpreted carefully.

**FIG. 1:**
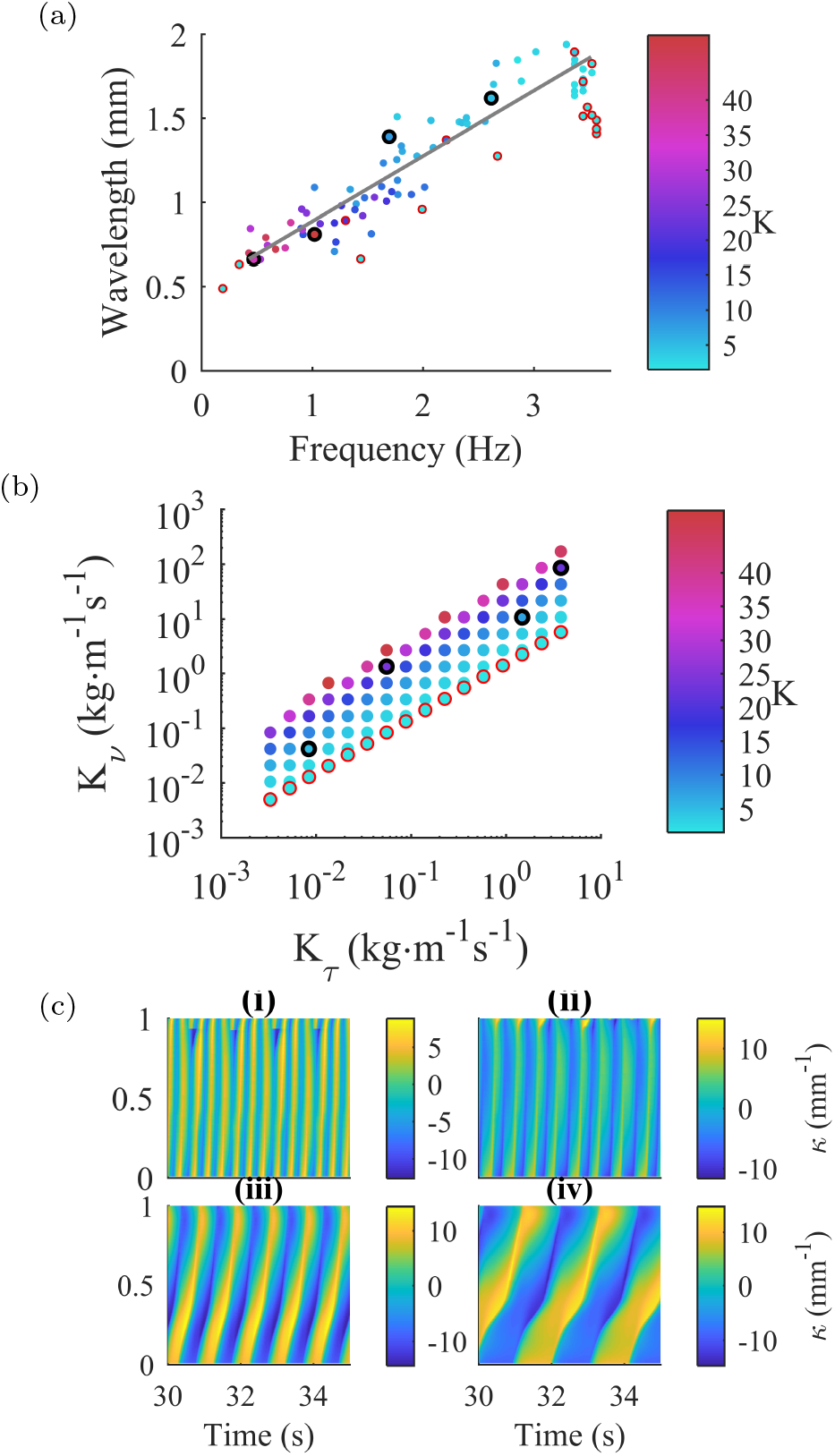
(a) Our model, with default parameters, reproduces frequency-wavelength relationship with increasing ratio, *K*, of tangential and normal drag coefficients *K_τ_* and *K_ν_*. (b) *K_τ_* and *K_ν_* values used in top panel, highlighting Newtonian environments (red circles) and parameters used for sample simulations (black circles). (c) Sample kymograms showing curvature dynamics in environments from water (i) to agar (iv).

It is worth noting that two models, presented here and in Boyle *et al*.^4^ have now demonstrated similar proprioceptively facilitated gait modulation, suggesting that these results emerge from the model assumptions rather that from implementation details (such as using continuous elastic shells versus articulated bodies).

## BODY ELASTICITY DICTATES THE ACCESSIBLE RANGE OF KINEMATIC PARAMETERS

The range of estimates for the Young’s modulus in *C. elegans* ranges over five orders of magnitude^23,26^ (from 3.77 kPa to 380 MPa). The methods used to obtain these estimates vary considerably and address complementary aspects of the worm’s material properties (see the discussion in Cohen and Ranner,^9^ for example). Importantly, biological muscle is an active material, and as such its Young’s modulus is itself modulated dynamically as a function of activity and internal state (length and speed), further complicating our limited knowledge of the passive material properties of the worm. We have previously considered the role of the Young’s modulus in facilitating undulations and forward thrust (or speed) in a mechanical framework driven by feed-forward (CPG-like) control.^9^Here we revisit the question in the context of a proprioceptively driven control. Whereas in a feed-forward setting, the question reduces to the ability of the body to follow the periodic driving force, within a closed loop control system the body shape is integral to the pattern generation. Thus, elasticity is expected to affect not only the response to the drive, but also the waveform and frequency of undulations, speed and neuromechanical phase lags.^11^ We therefore aim to characterize the kinematics of locomotion for different values of the Young’s modulus of the model.

We note that the mechanical model can be reformulated in nondimensional form.^9^ Nondimensionalization allows the physical parameters of a system to be recast in terms of fundamental control parameters of the system. In one such reformulation adopted here,^9^ the three parameters *K_ν_*, *K_τ_* and *E* can be related in terms of two nondimensional variables e and K expressed as

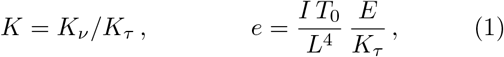

where *I* and *L* are constant geometrical factors representing the shape and length of the worm and *T*_0_ is a time constant. In this formulation, an increase in the elasticity *E* corresponds to holding *K* constant while increasing the value of e which this is equivalent to decreasing both tangential and normal environmental drag coefficients (*K_ν_*, *K_τ_*) such that their ratio remains constant, e.g., in Newtonian fluids (for *K* ≈ 1.5).

Simulation results (Fig. 2) confirm the above reasoning. The modulation of elasticity acts to modulate the gait, preserving the relation between anterior wavelength (see Appendix) and frequency obtained by modulating the environmental resistance, *K_τ_*. Interestingly, as the wavelength increases along the body, this relationship is violated.

**FIG. 2:**
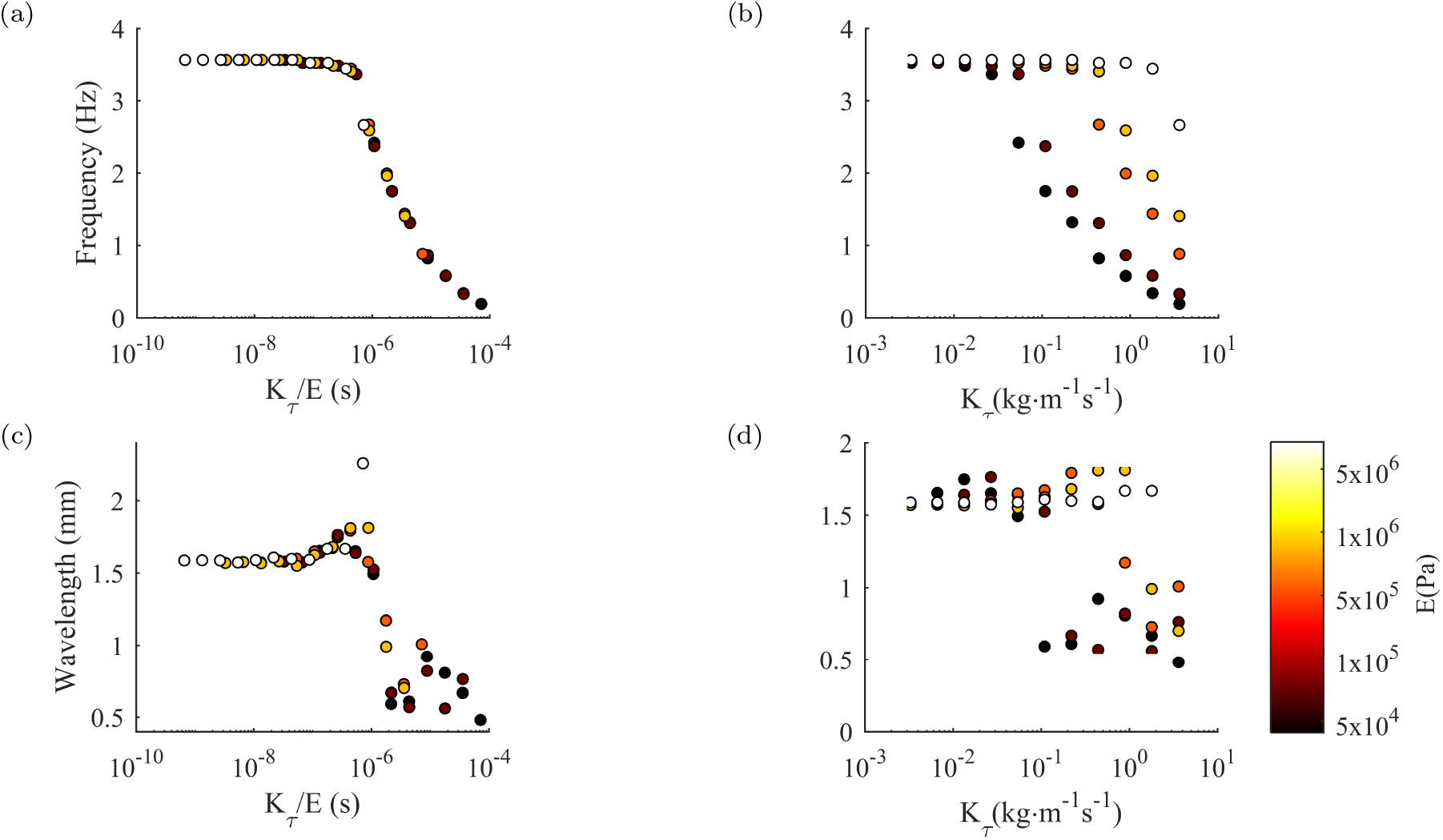
(a) Frequency over a range of *K_τ_* and *K* = 1.5 corresponding to Newtonian environments, simulated for different values of the Young’s modulus of the body. Higher elasticity limits the range of achievable frequencies and wavelengths within this range of Newtonian environments. Scaling by the Young’s modulus collapses data to a single relation. (b) The same data before scaling by Young’s modulus. (c) Anterior wavelength (but not wavelength) for the same simulations, scaled by the Young’s modulus also collapses to a single relation. (d) Anterior wavelength without scaling. Outliers represent simulations of worms exhibiting uncoordinated locomotion.

Within this purely elastic formulation of the body, we find two regimes: First, for a given choice of Young’s modulus, there exists a corresponding cutoff in *K_τ_*, below which the frequency and wavelength of undulations are saturated at their maximal (physiologically unrealistic) value. A higher Young’s modulus means that this cutoff is higher. It is in this low *K_τ_* that body damping is most important. Second, above the cutoff in *K_τ_*, increased fluid viscosity exerts growing mechanical load on the body, hence suppressing the frequency and wavelength of undulations. The higher the elasticity, the more the body resists this mechanical load. Within a given model (and parameters) of neuromuscular control, this effect can be used to estimate the effective elasticity of a worm that would be required to recapitulate the observed kinematics of the swim-crawl transition. Specifically, for the default parameters shown here, a Young’s modulus of the order of 50-100 KPa captures the full range of frequencies observed experimentally. In what follows, we therefore revert to our default parameter value of 100 KPa.

## PROPRIOCEPTIVE THRESHOLD INFLUENCES THE RANGE OF ACHIEVABLE GAIT

Like all animals, *C. elegans* is capable of adapting its gait in a context dependent manner, without changes in environmental viscosity. This must be achieved by some internal mechanism. Candidate mechanisms vary from descending neural control (e.g., a modulation of the current input from locomotion command neurons), a modulation of the motor circuit (e.g., the excitability of B-type motor neurons), a modulation of the proprioceptive field (though no such mechanism has been documented to date), etc. Although the neural model used here is idealized, we are able to vary the threshold value of the motor neurons. Here, an increased threshold could correspond to a reduced excitability of B-type motor neurons, reduced sensitivity to stretch, or a reduced tonic input current, e.g., from AVB neurons.

Fig. 3 shows the effect of threshold modulation in different environments. For threshold values below those shown in Fig. 3, undulations cease to occur, and for higher values the initial transient becomes very long (≳40 seconds). Our first observation is that undulations are robust for a wide range of thresholds. Furthermore gait modulation as a function of environmental viscoelasticity, or *K*, is robust to the choice of threshold. As expected, increasing the proprioceptive threshold (lowering sensitivity) results in a lower undulation frequency. However, unlike external modulation, the increased threshold manifests in an increased wavelength of undulation. Thus, the model predicts that the direct or indirect modulation of the proprioceptive threshold should lead to an inverse relationship between wavelength and frequency – as the frequency is increased, the wavelength should decrease and vice versa. Importantly, in the model, this inverse frequency-wavelength relationship is maintained across a wide range of environmental resistances, suggesting that the worm’s ability to modulate its waveform internally does not depend on environmental resistance (though the specific range of frequencies and wavelengths available vary in different environments).

**FIG. 3:**
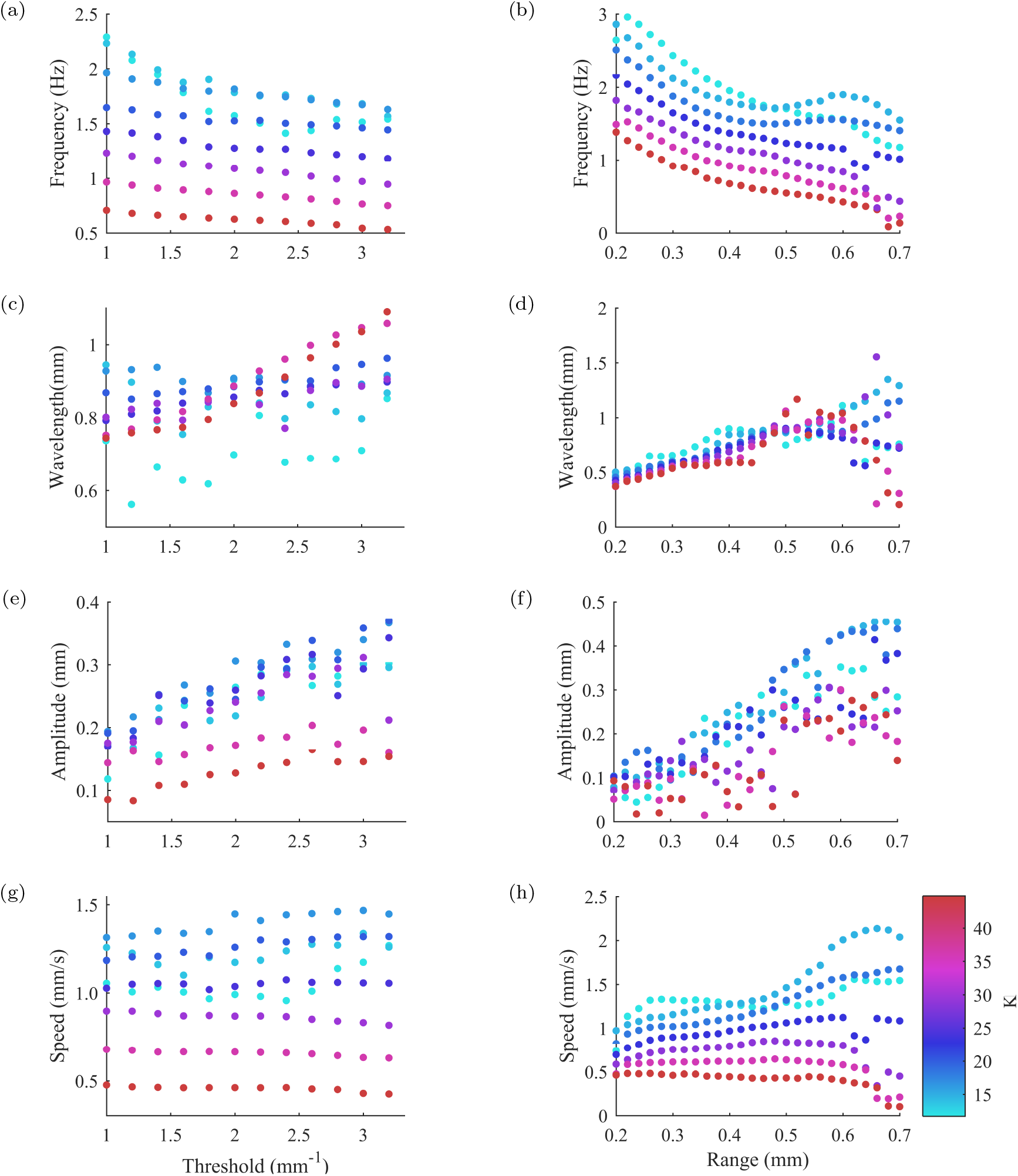
Internally modulated gait. Left: Proprioceptive threshold smoothly modulates locomotion kinematics over a range of model environments and threshold, switch strongest effects on amplitude and wavelength. Right: Proprioceptive range dramatically affects speed, undulation frequency and waveform. For both forms of internal modulation, note the decrease in undulation frequency as the wavelength and wave amplitude increase.

Overall, threshold modulation over the range tested appears to yield at most a 60% change in frequency and at most a 50% change in wavelength, as contrasted with the multi-fold range of frequencies and more than two fold increase in wavelengths arising from external gait modulation. As expected, the wave amplitude also increases with proprioceptive threshold; thus, as the frequency of undulations decreases, the wavelength increases and so does the wave amplitude. Finally, the speed of the worm in these simulations roughly corresponds to observed locomotion velocities across the entire range of thresholds tested.

## PROPRIOCEPTIVE RANGE IS THE MAIN DETERMINANT OF THE UNDULATION WAVELENGTH AND FREQUENCY IN A HOMOGENEOUS ENVIRONMENT

Proprioceptive control of undulations in *C. elegans* has long been postulated to rely on sensing of proximate and distal body or muscle length. Within the motor circuit of the ventral nerve cord, B-type motor neurons were postulated to fill this role through their extended posteriorly facing axons.^31^ The capacity to support locomotion through such a mechanism has since been confirmed in a series of modeling studies,^4,11,22^ but the only study to date to examine this in the context of swimming has required an anatomically unrealistic proprioceptive range of half a body length.^4^ To understand the role of sensory range in locomotion, and to better understand the link between sensory range and kinematics, we varied the proprioceptive range from a minimum of 20% to a maximum of 70% of the body length, posteriorly from the action of the muscle moment.

We find that increasing the range of proprioception in our model suppresses the frequency of undulations while extending the undulatory wavelength (Fig. 3). This effect is qualitatively the same as that of increasing the threshold, although the extent of the modulation of both frequency and wavelength is much larger. For sufficiently high external drag, the range of speeds achieved by simulated worms appears stable for all proprioceptive ranges, indicating that even local proprioception suffices to achieve robust locomotion. However, for lower viscoelasticity of the medium, speed is strongly affected by the proprioceptive range. These results are qualitatively consistent with those of Boyle *et al.*.^4^ Overall, it appears that within this model, each environment corresponds to a different ‘optimal’ proprioceptive range for maximizing locomotion speed. Surprisingly (but also consistent with Boyle *et al*.^4^), our simulation results indicate that a proprioceptive range of approximately half a body length best captures the experimentally observed locomotion metrics of frequencies, wavelengths, amplitudes and speed in different fluids across the swim-crawl transition.

## DISCUSSION

The importance of proprioception is well established in controlling posture and locomotion in a variety of limbless and legged species, from invertebrates to humans.^2,14,17,25,32^ And yet, the so-called sixth sense is least understood, from the molecular basis and biophysics of mechanosensation to the roles of proprioception in shaping motor patterns in peripheral and central nervous systems. In *C. elegans*, proprioception has been studied primarily with focus on the peripheral motor circuit, either with respect to posture, or to locomotion.^18,19,29^ Interestingly, most neurons associated with proprioceptive function contain axons that extend along the rostro-caudal body axis sublaterally or in the ventral nerve cord, suggesting an extended receptive field. Here, and in previous studies, we have followed this conjecture, although in other species, stretch receptors in neurons and muscles have been found to respond to deformation, muscle tension, or length.^32^ If stretch receptors integrate length along their body, it is essential to identify their receptive field in order to better understand the sensory motor loop.

While most conjectured proprioceptive neurons have posteriorly facing axons, Wen *et al*.^29^ have reported behaviors consistent with anteriorly facing proprioceptive fields, with a range of under 200 μm. Here, we showed (consistent with Boyle *et al*.^4^) that an extensive proprioceptive range (of approximately half the body length) is required in this model to generate the experimentally observed ranges of frequency and wavelength. Shorter ranges, while still generating robust locomotion, exhibit reduced wavelengths and a much reduced gait modulation. In particular, we see a saturation of the wavelength when increasing the proprioceptive range. This saturation occurs at shorter proprioceptive ranges (and shorter wavelengths) for increasingly resistive environments. Simulations using feedforward control of the same mechanical framework previously suggested^9^ that gait modulation may be needed by *C. elegans* to maximize its speed in different media. If so, the worm’s effective proprioceptive range places important constraints on the mechanical and sensory coupling mechanisms required for robust locomotion. That said, B-type neurons do not extend over half a body length, suggesting an important limitation in our understanding of this system. Our model therefore begs for a proposed mechanism by which such an effective proprioceptive range may be achieved in the worm.

We have compared the effects of mechanical load with those of two internal factors that likely affect the kinematics of forward locomotion. Our continuum neuromechanical model has reproduced gait adaptation and shown that two internal factors influence the frequency, wavelength and amplitude of undulation. In particular, we observe a qualitative distinction between mechanical and neural modulation. On the one hand, the model captures the positive correlation between frequency and wavelength as a function mechanical load.^1^ In contrast, in our model, increasing either the activation threshold or the proprioceptive field yields the opposite relationship: the higher the frequency, the lower the wavelength.

In this study, we have limited our consideration to a proprioceptive control mechanism, with a view to better understanding sensory-motor coupling effects subject to proprioceptive entrainment. To maximize the explanatory power of our investigation, we have simplified the sensory-motor coupling to a minimal model. This investigation therefore paves the road for further studies that may include a more detailed description of the neural circuitry and neuronal properties. In particular, we anticipate the fundamental insights gained to generalize to cases where such a proprioceptive mechanism is superimposed on centrally generated patterns, though this was not examined in detail in this work.

## ACKNOWLEDGMENTS

We thank Felix Salfelder for useful discussions. This work was funded by the EPSRC (EP/J004057/1). TR is funded by a Leverhulme Trust Early Career Fellowship.

## APPENDIX COMPUTING KINEMATIC PARAMETERS

All simulations were performed for 60 seconds using integration time steps of 0.3 ms. Some transient durations were negligible (<1 second) but others varied significantly with model parameters. In all kinematic analysis, we truncated the transient, thus limiting our analysis to periodic activity.

### Frequency

The period of undulations, *T*, was computed from curvature kymograms. For a given point along the body, the period was defined as the mean time interval between zero crossing of the body curvature *ν*(from negative to positive values). In coordinated locomotion, the period of undulation does not depend on the position along the body. The frequency of undulations is given by *f* =1/*T*.

### Wavelength

The computation of wavelength was more complicated since, in our proprioceptive model (as in experimental observations), the wavelength increases along the body (from head to tail). We defined wavelength as the distance along the body spanning an entire cycle of body curvatures^1^. We computed wavelength by calculating the gradient of the curvature k as a function of body coordinate u and time t within a section of the body. We use two different sections along the body, defining *wavelength λ* over *u* ∈ (0.1, 2/3) mm and *anterior wavelength λ*_ant_ over *u* ∈ (0.1,0.3) mm. The corresponding wavelength was then given by the product of the period and gradient 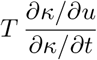. A small amount of filtering ensured that the derivatives are well approximated using a finite difference. An average was taken across all available space *u* and time *t* using a histogram mode with logarithmically distributed bins. In this paper, the wavelength *λ* was used in all figures except in our analysis of elasticity (Fig. 2) and proprioceptive range (Fig. 3) which show the anterior wavelength instead.

### Undulation amplitude

Undulation amplitude also varies along the body. Here it was computed by measuring the perpendicular distance of a single body coordinate, *u* = 0.4, along body to the line connecting the head and tail. A peak amplitude was obtained in every undulation period, and peak amplitudes were averaged over time, yielding a mean undulation amplitude.

### Speed

Speed was computed by tracking the midpoint of the worm’s body (*u* = 0.5) over time, and performing a straight line fit over the midpoint trajectory to remove side-to-side displacement arising from the undulatory movement. The speed was then defined as the distance traveled along the straight line over the corresponding time interval.

## References

1 S. Berri, J.H. Tassieri. M. Boyle, I.A. Hope, and N. Cohen. Forward locomotion of the nematode c. elegans is achieved through modulation of a single gait. HFSP Journal, 3(3):186–193, 2009. PMID: 19639043.

2 R. Blake and R. Sekuler. Perception. McGraw-Hill, NY, NewYork, 5 edition, 2005.

3 J. Boyle. C. elegans locomotion: An integrated approach. unpublished thesis, 2009.

4 J.H. Boyle, S. Berri, and N. Cohen. Gait modulation in c. elegans: an integrated neuromechanical model. Frontiers in computational neuroscience, 6:10, 2012.

5 J.A. Bryden and N. Cohen. Neural control of caenorhabditis elegans forward locomotion: the role of sensory feedback. Biological cybernetics, 98(4):339–351, 2008.

6 I. Buntschuh, D.A. Raps, I. Joseph, C. Reid, A. Chait, R. Totanes, M. Sawh, and C. Li. Flp-1 neuropeptides modulate sensory and motor circuits in the nematode caenorhabditis elegans. PloS one, 13(1):e0189320, 2018.

7 M. Chal?e, J.E. Sulston, J.G. White, E. Southgate, J.N. Thomson, and S. Brenner. The neural circuit for touch sensitivity in caenorhabditis elegans. J. Neurosci., 5(4):956–964, 1985.

8 N. Cohen and J. H. Boyle. Swimming at low reynolds numbers: a beginners guide to undulate ry locomotion. Contemp. Phys., 51:1–21, 2010.

9 N. Cohen and T. Ranner. A new computational method for a model of C. elegans biomechanics: Insights into elasticity and locomotion performance. ArXiv e-prints, Feb 2017.

10 N. Cohen and T. Sanders. Nematode locomotion: dissecting the neuronal–environmental loop. Current opinion in neurobiology, 25:99–106, 2014.

11 J.E. Denham, T. Ranner, and Cohen. N. Neuromechanical phase lag predicts material and control properties in *Ceanorhabditis elegans*. Phil. Trans. R. Soc. Lond. B, Biol. Sci., 2018.

12 C. Fang-Yen, M. Wyart, J. Xie, R. Kawai, T. Kodger, S. Chen, Q. Wen, and A.D.T. Samuel. Biomechanical analysis of gait adaptation in the nematode caenorhabditis elegans. Proc. Nat. Acad. Sci. U.S.A, 107(47):20323–20328, 2010.

13 C. Fieseler, J. Kunert-Graf, and N.J. Kutz. The control structure of the nematode caenorhabditis elegans: neuro-sensory integration and propioceptive feedback. arXiv preprint arXiv:1707.05359, 2017.

14 E. Fuchs, P. Holmes, David I., and Ayali A. Proprioceptive feedback reinforces centrally generated stepping patterns in the cockroach. J. Exp. Biol., 215:1884–1891, 2012.

15 J. Gray and H.W. Lissmann. The locomotion of nematodes. J. Exp. Biol., 41(1):135–154, 1964.

16 Z.V. Guo and L. Mahadevan. Limbless undulatory propulsion on land. Proc. Nat. Acad. Sci. U.S.A., 105(9):3179–3184, 2008.

17 A.A.V. Hill, M.A. Masino, and R.L. Calabrese Calabrese. Intersegmental coordination of rhythmic motor patterns. J. Neurophysiol., 90(2):531–538, 2003. PMID: 12904484.

18 W. Li, Z. Feng, P.W. Sternberg, and X.Z.S. Xu. A c. elegans stretch receptor neuron revealed by a mechanosensitive TRP channel homologue. Nature, 440:684–687, 2006.

19 X. Liang, X. Dong, D.G. Moerman, K. Shen, and Wang X. Sarcomeres pattern proprioceptive sensory dendritic endings through perlecan/unc-52 in c. elegans. Dev. Cell, 33(4):388–400, 2015.

20 J. Lighthill. Flagellar hydrodynamics. SIAM review, 18(2):161–230, 1976.

21 S. L. McIntire, E. Jorgensen, J. Kaplan, and H. R. Horvitz. The GABAergic nervous system of caenorhabditis elegans. Nature, 364:337–341, 1993.

22 E. Niebur and P. Erdös. Theory of the locomotion of nematodes: Dynamics of undulatory progression on a surface. Biophysical Journal, 60(5):1132–1146, 1991.

23 S-J Park, M.B. Goodman, and B.L. Pruitt. Analysis of nematode mechanics by piezoresistive displacement clamp. Proc. Nat. Acad. Sci., U.S.A, 104(44):17376–17381, 2007.

24 B.C. Petzold, S-J Park, P. Ponce, C. Roozeboom, C. Powell, M.B. Goodman, and B.L. Pruitt. Caenorhabditis elegans body mechanics are regulated by body wall muscle tone. Biophys. J., 100(8):1977–1985, 2011.

25 C.S. Sherrington. The integrative action of the nervous system. Cambridge University Press, Cambridge, 1906.

26 J. Sznitman, P.K. Prashant, P. Krajacic, T. Lamitina, and P.E. Arratia. Material properties of caenorhabditis elegans swimming at low reynolds number. Biophys. J., 98(4):617–626, 2010.

27 A. Vidal-Gadea, S. Topper, L. Young, A. Crisp, L. Kressin, E. Elbel, T. Maples, M. Brauner, K. Erbguth, A. Axelrod, et al. Caenorhabditis elegans selects distinct crawling and swimming gaits via dopamine and serotonin. Proc. Nat. Acad. Sci. U.S.A, 108(42):17504–17509, 2011.

28 H.R. Wallace. Wave formation by infective larvae of the plant parasitic nematode meloidogyne javanica. Nematologica, 15:65–75, 1969.

29 Q. Wen, M.D. Po, E. Hulme, S. Chen, X. Liu, S.W. Kwok, M. Gershow, A.M Leifer, V. Butler, C. Fang-Yen, T. Kawano, W.R. Schafer, G. Whitesides, M. Wyart, D.B. Chklovskii, M. Zhen, and A.D. Samuel. Proprioceptive coupling within motor neurons drives c. elegans forward locomotion. Neuron, 76(4):750–761, 2012.

30 J.G. White, E. Southgate, and J. Nichol Thomson. The structure of the ventral nerve cord of caenorhabditis elegans. Phil. Trans. R. Soc. Lond. B, 275(938):327–348, 1976.

31 J.G. White, E. Southgate, J.N. Thomson, and S. Brenner. The structure of the nervous system of the nematode caenorhabditis elegans. Philos. Trans. R. Soc. London B, Biol. Sci., 314:1–340, 1986.

32 X. Yu, B. Nguyen, and O. W. Friesen. Sensory feedback can coordinate the swimming activity of the leech. J. Neurosci., 19(11):4634–4643, 1999.

